# Amyloid damage to islet β-cells in type 2 diabetes: hypoxia or pseudo-hypoxia?

**DOI:** 10.1101/810747

**Authors:** Vittorio Bellotti, Alessandra Corazza, Beatrice Foglia, Erica Novo, J. Paul Simons, P. Patrizia Mangione, Guglielmo Verona, Diana Canetti, Paola Nocerino, Laura Obici, Alessandro Vanoli, Marco Paulli, Sofia Giorgetti, Sara Raimondi, Raya Al-Shawi, Glenys A. Tennent, Graham W. Taylor, Julian D. Gillmore, Mark B. Pepys, Maurizio Parola

## Abstract

Aggregation of islet amyloid polypeptide (IAPP) and amyloid deposition in the islets of Langerhans may significantly contribute to the multifactorial pathogenic mechanisms leading to type 2 diabetes. A direct toxic effect on β-cells of oligomeric IAAP has been demonstrated in *in vitro* models, but the mechanism operating *in vivo* is still unclear. Mice models presenting amyloid deposition and glucose intolerance represent a good tool for exploring *in vivo* a putative mechanism of toxicity directly related to the physical expansion of the extracellular matrix by the amyloid fibrillar aggregates. Based on our hypothesis that deposition of amyloid may influence the oxygen perfusion, we have calculated that the mean distribution of oxygen partial pressure would drop by more than 50 % in the presence of amyloid deposits in the islet. This condition of hypoxia caused by the remodelling of the extracellular space may explain the metabolic abnormalities in the Langerhans islets, otherwise interpreted as pseudo-hypoxic response to IAPP oligomers.

---

Montemurro et al.^1^ have recently shown that *in vitro* exposure to oligomeric human islet amyloid polypeptide (IAPP) triggers the hypoxia response in islets of Langerhans from diabetic IAPP transgenic rats. In the absence of actual hypoxia, this response was considered pseudo-hypoxic. The metabolic shift of β-cells toward aerobic glycolysis and depression of Krebs cycle in type 2 diabetes, attributed to mitochondrial toxicity of IAPP oligomers, was suggested to resemble the Warburg effect in tumours^1,2^. There is also transcriptomic and proteomic evidence of hypoxia in islets of Langerhans in type 2 diabetes.

The islets of Langerhans of type 2 diabetes patients^3^ and human islets that fail after heterotopic transplantation^4^ usually contain typical amyloid deposits composed of insoluble fibrillar IAPP. However, the relative pathogenic role of different amyloid protein conformers, especially soluble oligomers and insoluble amyloid fibrils, in actually damaging cellular and tissue function *in vivo* has not been established. *In vitro* islet models have all focussed on soluble oligomeric aggregates but defining the effects of different ‘amyloid’ conformers in type 2 diabetes is very challenging. The pathogenesis of type 2 diabetes is multi-factorial and studying cellular effects of actual fibrillar amyloid deposits *in vitro* is particularly intractable. We have therefore investigated potential pathogenic mechanisms *in vivo* in a human IAPP transgenic mouse type 2 diabetes model. Here we show that significant expansion of the extracellular space by islet amyloid deposits increases the distance between the capillary lumen and the β–cells. This unequivocal histopathological effect of fibrillar amyloid deposition on the islet architecture must inevitably impair the oxygen pressure gradient, causing actual hypoxia for the β–cells.

## Results and Discussion

A^vy^ mice^5^ spontaneously become diabetic and the phenotype is exacerbated by transgenic expression of human IAPP (Fig. 1), associated with progressive accumulation of human IAPP amyloid in their islets of Langerhans from about 4 months of age (Fig. 2). The amyloid is deposited throughout the islet extracellular space, within the capillary walls and expanding the distance between the capillary endothelium and the endocrine secretory cells (Fig. 2C). Only traces of hypoxia inducible factor (HIF)-1α and HIF-2α subunits were detected in amyloid-free islets of control, age matched A^vy^ mice (Fig. 3A), but the β–cells of diabetic mice with islet amyloid deposition overexpressed both HIF-1α and HIF-2α (Fig. 3B). Our *in vivo* observations thus confirm the molecular signs of hypoxia reported in the islets of human IAPP transgenic rats exposed to human IAPP *in vitro*^1^.

**Figure 1.**
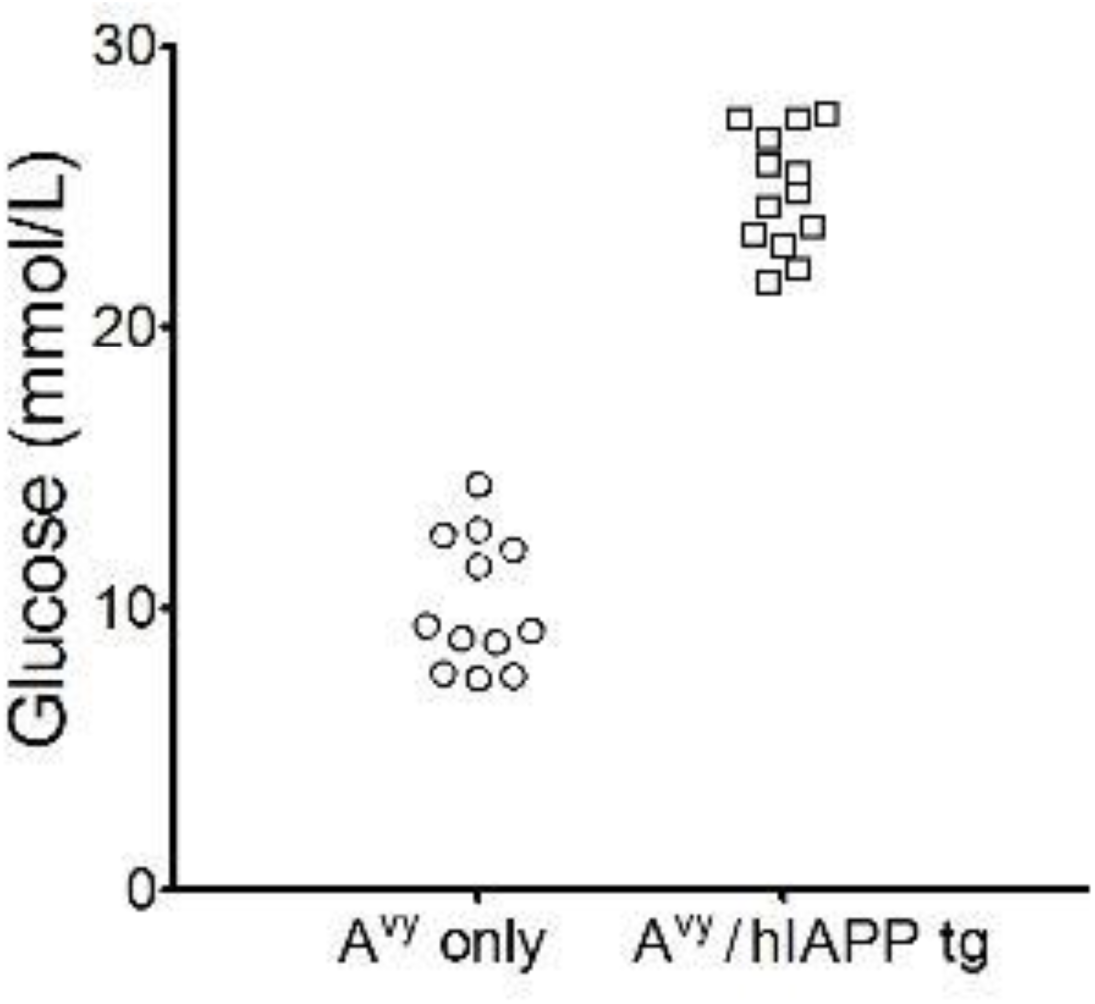
Hyperglycaemia in human IAPP transgenic A^vy^ mice. Glucose concentrations 60 min after standard glucose challenge in individual male control A^vy^ mice (*n*=12) and hIAPP transgenic A^vy^ mice (*n*=13).

**Figure 2.**
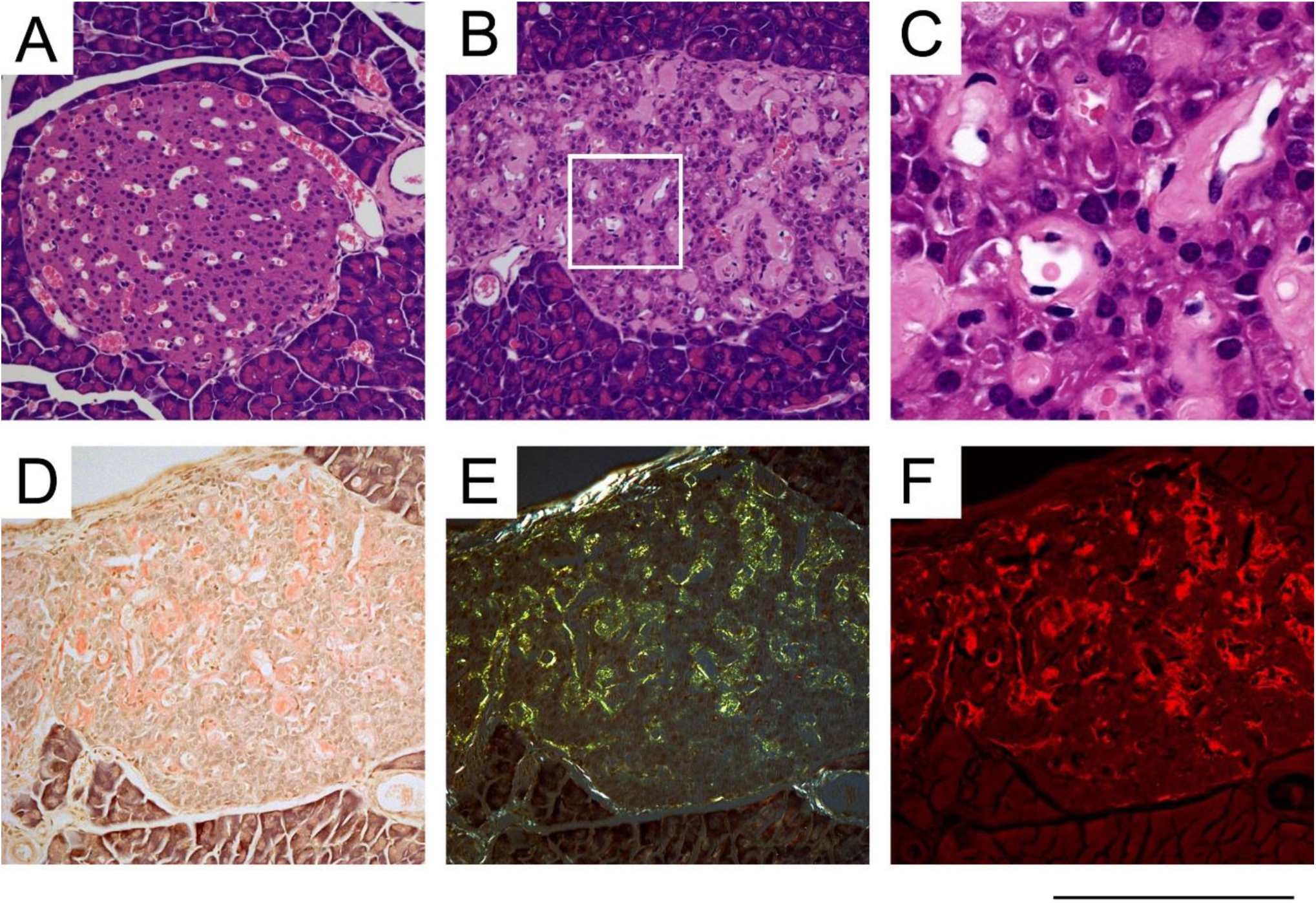
Islet amyloid deposition in human IAPP transgenic A^vy^ mice. A: Typical pancreatic islet from a control 7-month old, non-transgenic, male, control A^vy^ mouse. B-F: Pancreatic islets from a 7-month old, male, hIAPP transgenic A^vy^ mouse with abundant amyloid. Amyloid deposits are seen as homogeneous, eosinophilic, Congo red-positive extracellular layers between islet endocrine cells and capillaries. A-C: H&E stain; D-F: Alkaline alcoholic Congo red stain of 6 μm sections^13^. D: bright field viewed with 10x objective, showing the amyloid load. E: intense cross polarised illumination at full extinction showing the pathognomonic green birefringence of amyloid. F: UV fluorescence microscopy showing sensitive detection of amyloid. Scale bar: 250 µm (A, B, D, E, F); 65 µm (C).

**Figure 3.**
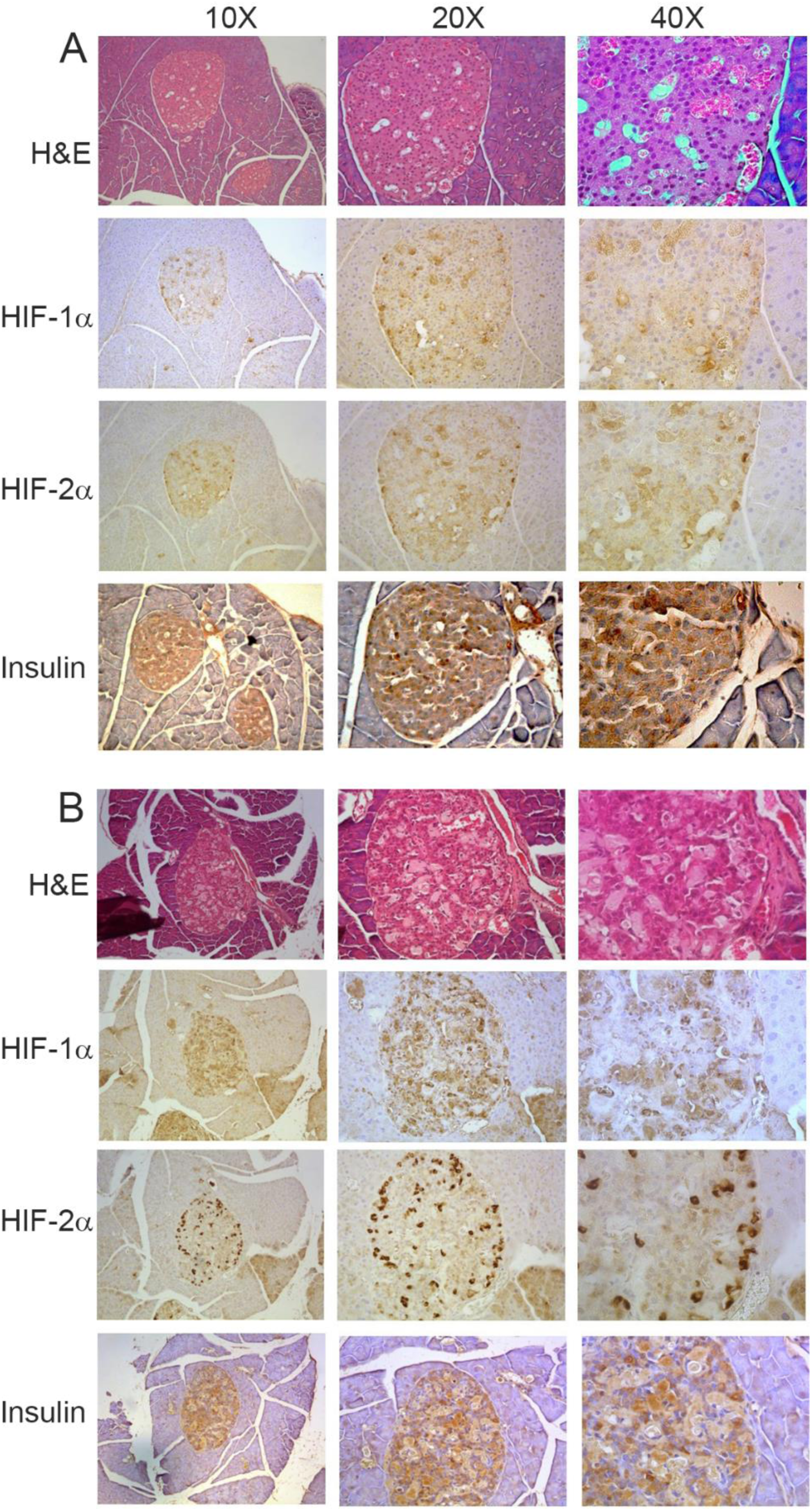
Increased β-cell expression of hypoxia inducible factors (HIFs) in amyloidotic pancreatic islets. Formalin fixed pancreas from (A), a 7-month old control, non-transgenic, male A^vy^mouse and (B), a 7-month old, human IAPP transgenic, male A^vy^ mouse with abundant islet amyloid. Two µm sections were stained with haematoxylin and eosin (H&E). Four µm sections were immunoperoxidase stained for HIF-1α, HIF-2α and Insulin as previously described^14,15^. β-cells identified by positive staining for Insulin and by their morphology. They were clearly positive for HIF-1α and, more strongly, for HIF-2α, in the amyloidotic but not control islets. Specificity of the immunostaining was confirmed by consistently negative controls, including isotype and concentration matched irrelevant antibody in place of the primary antibodies.

The oxygen supply of islet cells depends on various factors, including arterial oxygen partial pressure, vascularization within the pancreas, diffusion of oxygen across capillary walls and across the extravascular, extracellular space. In order to elucidate the pathogenesis of the islet hypoxia observed *in vivo*, we therefore measured the distances between capillaries and β-cells in our control and amyloidotic mice, as shown in Figure 4A. The frequency distributions of 293 values from 6 individual control islets and 448 values from 24 individual amyloidotic islets (Fig. 4B) had means ± SD of 5.8 ± 2.6 µm and 14.3 ± 6.4 µm respectively. The islet amyloid deposits clearly increase the space across which oxygen must diffuse, between the capillary wall and the β–cells. We quantified this effect by modelling the decrease of oxygen partial pressure, PO2, across the extracellular space by solving the steady state reaction-diffusion equation^6^

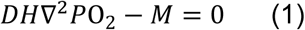

where *D* is the oxygen diffusion coefficient, *H* is the oxygen solubility coefficient and *M* is the rate of irreversible chemical reaction using O_2_. If we assume P_0_ = 100 mmHg, just outside the capillary wall, it follows, according to the simple diffusion model for homogeneous tissues, that the oxygen can penetrate into the tissues for a distance 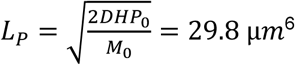, and that after this limit value the oxygen tension is zero. The profile of the oxygen tension along the tissue (Fig. 4C) shows that PO2 drops from 65 to 27 mmHg for controls and amyloidotic tissues.

**Figure 4.**
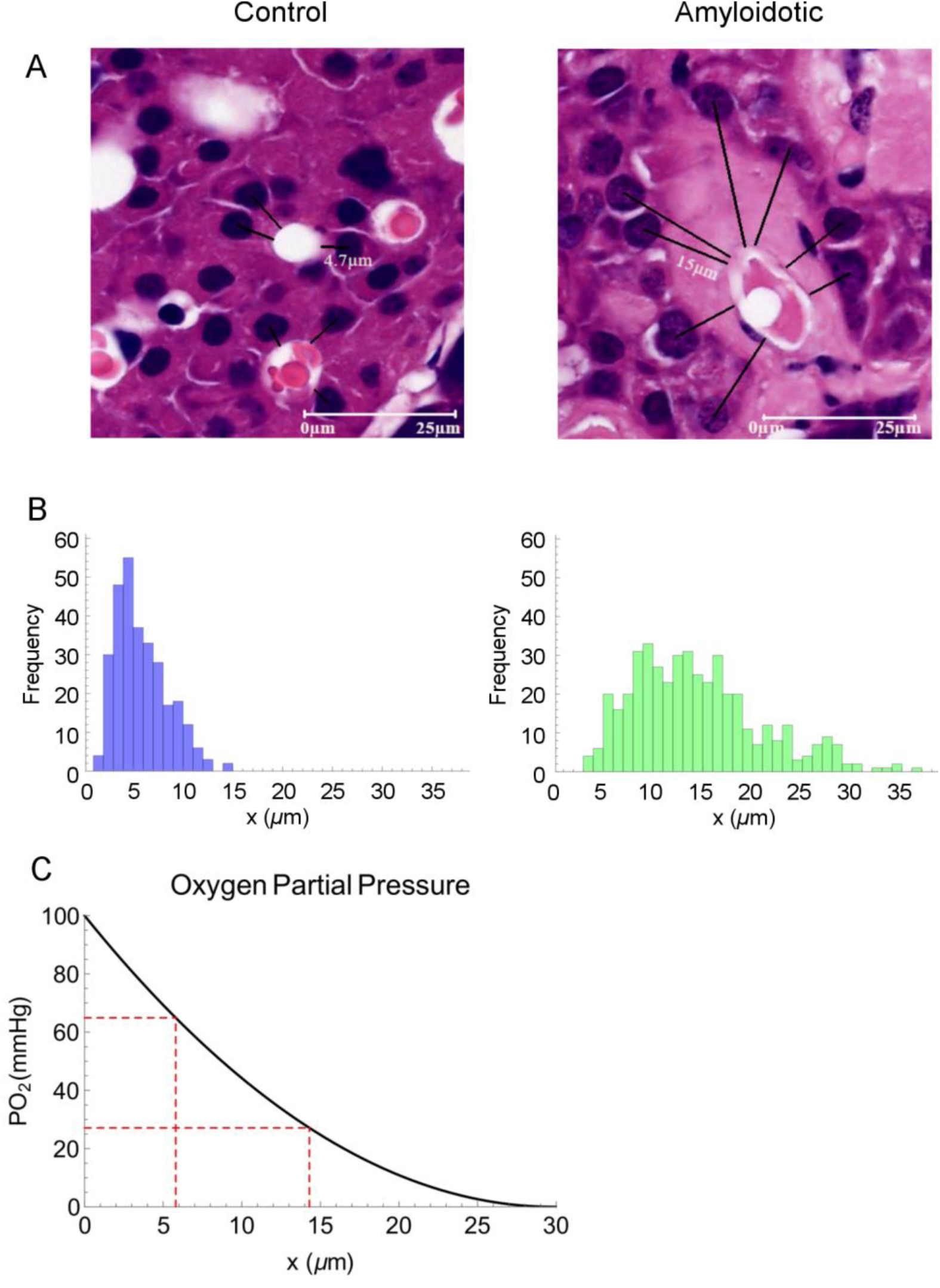
Extended capillary-cell distances in amyloidotic islets reduce the PO2 at the β-cells. (A). Black lines in the images of control and amyloidotic islets show the distances from the capillary wall to the centre of the nucleus of surrounding β-cells. These distances were measured directly in mm on each image by GIMP image editor software and then converted to µm using the 25 µm scale bar (solid white line) derived from calibration of the microscope objective. (B). Frequency distribution of distances showing a highly significant difference between the average in control and amyloidotic tissues, *t-*test p-value = 8.4 × 10^-82^. The PO_2_ values across the distance from capillary to cells (C) were calculated by solving equation 1. The PO_2_ values at 5.8 and 14.3 µm were 65 and 27 mmHg for normal and amyloidotic tissues, respectively.

The diffusion coefficient used to determine the PO_2_ curve (Fig. 4C) was that reported for animal pancreatic islets^7^. It is not known for amyloidotic tissues therefore we estimated the PO_2_ variation in different diffusion conditions using limiting values of D reported for other tissues, ranging from 1 to 4 × 10^-9^ [m^2^/s]^8^. With these constraints, the range of L_P_ is 26-52 µm and the PO_2_ difference between normal and amyloidotic tissues ranges from 40-26 mmHg. Thus, even if the actual value of the diffusion coefficient within the pancreas differs from the one we have adopted, the phenomenon remains substantial and is biologically significant.

In conclusion, the β-cells of the islet of Langerhans, both in experimental and clinical type 2 diabetes, express the molecular signature of hypoxia. The signature is also seen when islets are exposed to amyloidogenic human IAPP *in vitro*, although it is not clear whether this reflects actual hypoxia or direct mitochondrial toxicity of IAPP oligomers. In contrast, our present findings highlight the incontrovertible role of the extracellular islet amyloid deposits. By expanding the interstitial space of the islets, these deposits inevitably compromise diffusion of oxygen from the capillary circulation. The synthesis and secretion of insulin by β-cells involves high oxygen consumption under normal physiological conditions^9^. Further damage by IAPP oligomers to β-cells, which are already struggling in a hypoxic environment, cannot be excluded. However, the parenchymal cellular hypoxia, inevitably caused by the deposition and progressive accumulation of amyloid in the extracellular matrix, is likely to be crucial. Indeed, such hypoxia may be a general mechanism by which amyloid of any type causes tissue damage, with particular importance in tissues with the highest oxygen consumption.

## Methods

### Mouse model of islet amyloidosis

Under normal conditions, hemizygous hIAPP transgenic mice express human IAPP, but are normoglycemic and do not accumulate IAPP amyloid^10^. Mice heterozygous for the spontaneous agouti viable yellow mutation (A^vy^) have yellow coats and become obese and diabetic, but they lack IAPP amyloid^11^. Mice heterozygous for A^vy^ and hemizygous for the hIAPP transgene exhibit a phenotype reminiscent of human type-2 diabetes, with accumulation of IAPP amyloid in islets of Langerhans^5^. Such mice were generated by crossing transgenic mice that express human IAPP in β-cells (Jax stock No: 008232) with agouti viable yellow mice (Jax stock No: 000017). To maximise the phenotypic expression of the A^vy^ allele (which is variably expressed), breeding was preferentially done using female mice with completely yellow coats^12^. The A^vy^/hIAPP mice used for experiment were completely or nearly completely yellow. The controls were A^vy^ mice (i.e. lacking the hIAPP transgene).

### Glucose tolerance test

Food was withdrawn for 6 h, following which the mice were challenged with 10 µl/g of 20 % glucose solution injected intraperitoneally. Blood glucose concentration was assayed 60 min later with a calibrated standard hand-held monitor (ACCU-CHEK Inform II).

### Histology

Wax tissue fixed overnight in neutral buffered formalin were analysed. Staining for amyloid was performed on 6 µm thick sections by the alcoholic alkaline Congo red method of Puchtler et al^13^. Different thickness was used for sections stained with haematoxylin and eosin (H&E, 2 µm) and for those used for immunohistochemistry (4 µm).

### Immunohistochemistry analysis

Paraffin pancreas sections of specimens derived from amyloidotic mice or related controls were used. Tissues were mounted on poly-L-lysine coated slides, were incubated with the polyclonal antibody against HIF-1α (Novus Biological Centennial, CO, USA dilution 1:150) or with the polyclonal antibody against HIF-2α (Novus Biological Centennial, CO, USA dilution 1:100), or with polyclonal antibody against insulin (Thermo Fisher Scientific, Rockford, IL, USA, dilution 1:100). After blocking endogenous peroxidase activity with 3 % hydrogen peroxide and performing microwave antigen retrieval (citrate buffer, pH 6), primary antibodies for HIF-1α and HIF-2α were labeled by using EnVision, HRP-labeled System (DAKO) and visualized by 3’-diaminobenzidine substrate^14^. For detection of insulin, as a marker of the cells, we used guinea pig antibodies (DAKO) according to the procedure described by Westermark and coauthors^15^. Negative controls were performed by replacing the respective primary antibodies by isotype and concentrations matched irrelevant antibody.

### Measurement of the distance between capillary and β-cells in pancreatic islets

To determine whether deposition of amyloid may influence the oxygen perfusion of the islets, H&E images of pancreatic specimens from amyloidotic and non amylodotic mice were acquired. Distances between the capillary walls to the centre of the nucleus of surrounding β-cells were measured directly on each image in mm using GIMP editor software and then converted to µm using the 25 µm scale bar of the microscope.

### Oxygen partial pressure determination

The oxygen diffusion in the tissue was estimated by solving the steady state reaction-diffusion equation

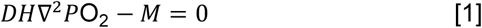

where *D*= 1.31 × 10^-9^ [m^2^/s]^7^ is the oxygen diffusion coefficient, *H*= 0.17 [mol O_2_/atm m^3^]^16^ is the oxygen solubility coefficient and *M* is the rate of irreversible chemical reaction using O_2_. In our model we have assumed the oxygen consumption rate constant M= M_0_ = ρκ, where the cell density ρ = 8.6 × 10^14^ [cells/m^3^]^17^ with an oxygen consumption rate κ = 7.6 × 10^-17^[mol O_2_/cells]^17^. The solution of equation 1 in the case of the simple geometry of a semi-infinite plane, for x > 0, with the boundary condition P = P_0_ at x= 0^6^is

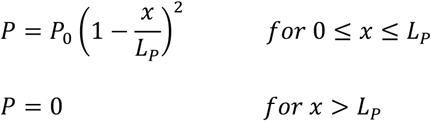

where 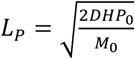

## Acknowledgements

This work was supported by grants from the U.K. Medical Research Council (MR/R016984/1), the Italian Ministry of Health (Ricerca Finalizzata RF 2013 02355259) and the Istituto Nazionale di Biostrutture e Biosistemi. Core support for the Wolfson Drug Discovery Unit is provided by the UK National Institute for Health Research Biomedical Research Centre and Unit Funding Scheme via the UCLH/UCL Biomedical Research Centre and by the UCL Amyloidosis Research Fund.

## Author contributions

The study was conceived, designed and supervised by V.B. and M.P. B.F., E.N., J.P.S., G.V., D.C., P.N., A.V., S.G., S.R., R.A-S., G.A.T. performed research. A.C., P.P.M., L.O., M.P., G.W.T., J.D.G. and M.B.P. contributed to experimental design and discussion. All the authors analysed and interpreted the data. The paper was written by V.B., A.C., M.P., and M.B.P. and reviewed and approved by all co-authors.

